# Impact of urbanization on the taxonomic and functional diversity of spider assemblages in Guwahati City, Assam, India

**DOI:** 10.1101/2023.01.07.523076

**Authors:** Ankita Sharma, Bitupan Deka, Puja Bishaya, Raman Kumar, Narayan Sharma

**Author notes:** Corresponding author: Narayan Sharma.

## Abstract

The homogenous nature of the urban environment rapidly alters community dynamics of extant flora and fauna due to short-term spatial and temporal factors. However, such impacts of urbanization are mostly investigated in terms of taxonomic diversity, while its impact on functional diversity remains poorly understood. Whereas taxonomic information is limited to the identity of species, functional traits determine the relationship between species identity and ecosystem functioning. Studies investigating the role of urbanization in altering these ecological parameters have mostly focused on avian communities or plant species, while arthropods such as spiders which are integral components of urban households have largely been overlooked. This study aims to understand the impact of urbanization on both taxonomic diversity and functional diversity of spider assemblages across an urban–semi-urban–forest gradient in Guwahati, a rapidly growing city in northeast India. We surveyed spiders at 13 sites representing four habitat types (urban, urban parks, semi-urban, and forests) using belt transects, and also recorded functional traits relating to key life history processes. Spider species composition differed significantly between various habitats. The taxonomic diversity of spiders was highest in forests and lowest in urban parks. The turnover component was the major contributor to changing the β-diversity of spiders. Reduced diversity in urban regions was likely due to the dominance of a few synanthropic species. Generalised linear mixed-effects model analysis indicated that the habitat types significantly impacted spider abundance. Functional richness was maximum in forests (F_ric_ = 23.43) and minimum in urban habitats (F_ric_ = 12.98), while functional divergence was maximum in urban sites (F_div_ = 0.79). Our study demonstrates that urban land-use change can alter the structure and functioning of the spider community.

## Introduction

More than half of the world’s population lives in urban areas – increasingly so in densely populated cities (Ritchie and Roser 2018); by 2050 it is projected to increase to 68% (UN DESA 2019). Deforestation, fragmentation and loss of connectivity caused by the rapidly increasing urbanisation have caused many wildlife species to live in close proximity to urban areas (Bonte et al. 2004; Gopalan and Radhakrishna 2022). Urbanization-induced habitat destruction and fragmentation are often the leading causes of changes in community structure and subsequent deterioration of ecological interactions between species and trophic systems (Seto et al. 2012; Lowe et al. 2018; Seibold et al. 2019). It is well-documented that urbanization leads to an overall loss of biodiversity (McKinney 2006; Evans et al. 2009; Theodorou 2022). For instance, the loss of avian species in various continents has been attributed to a lack of appropriate adaptations in most species for exploiting resources and avoiding risks in urban environments (Sol et al. 2014). Urbanization can thus restructure the biotic communities in urban areas due to the decline of species intolerant to urban development or, in some cases, an increase in abundance of more tolerant species (Hagen et al. 2017). A growing body of research has documented the negative impact of urbanization on the taxonomic diversity of various organisms including invertebrates (Buczkowski and Richmond 2012; Fenoglio et al. 2020; Piano et al. 2020), amphibians (Scheffers and Paszkowski 2012) and birds (Marzluff 2001; Horváth et al. 2012; Batáry 2017). Further, increasing urbanization also alters the β-diversity (Socolar et al. 2016; Tóth and Hornung 2020).

While many studies have assessed the impact of urbanization on taxonomic diversity, there is little information about the alteration in the functional traits of different organisms within an ecosystem in urban species assemblages. Functional diversity is the variation in the functional traits of different organisms that influence ecosystem functioning (Tilman 2001; Petchey and Gaston 2006). It is numerically represented using functional diversity indices that allow the quantification and comparison of functional diversity among communities (Pla et al. 2012). Despite the gap in knowledge concerning functional diversity in urban habitats, a few studies have shown that changes in the environment due to anthropogenic activities can negatively influence the functional diversity at the local scale (Sol et al. 2020). For example, the functional traits of spiders, birds, and millipedes are negatively influenced by urbanization (Croci et al. 2008; Meffert and Dziock 2013; Sol et al. 2014; Sánchez-Ruiz and Brescovit 2018; Toth and Hornung 2020). Moreover, urbanized systems exhibit higher rates of change in life-history traits than do natural habitats, indicating that urbanization can lead to evolutionary changes in the functional role played by a species in a community (Alberti et al. 2017). Functional diversity can serve as a biodiversity indicator because species represent a varying range of functional traits. Any changes in these traits are useful in understanding their sensitivity to ecosystem changes (Vandewalle et al. 2010).

Arthropods constitute a diverse group of organisms that provide a range of ecosystem services including pollination, nutrient cycling, biological pest control and decomposition (Fenoglio et al. 2020). Arthropods are characteristically sensitive to environmental changes and respond rapidly to them. Moreover, as arthropods can be surveyed relatively easily and economically, they serve as suitable indicators of environmental quality (Wettstein and Schmid 2001; Uehara-Prado et al. 2009). The impacts of urbanization on arthropods are mediated via pollution, draining and diversion of watercourses, and habitat loss or fragmentation (McIntyre 2000). The urban expansion also impacts plant–arthropod interactions and reshapes the abundance and behaviour of arthropods on urban plants (Dale and Frank 2018). Additionally, evidence suggests that urbanization exerts an overall negative impact on the richness of arthropod communities, resulting in their decreased diversity and abundance in urban landscapes (Martinson and Raupp 2013; Fenoglio et al. 2020). In India, there is limited research on the impact of urbanization on biodiversity. The few studies that have examined the role of urbanization in altering biodiversity patterns have focused on only limited taxa such as birds and larger mammals. Invertebrates, despite being a major contributor to regional biodiversity, are the least studied. Moreover, functional diversity has only recently begun receiving attention as an integral concept towards understanding community dynamics. How the impacts on functional diversity vary across ecosystems (i.e., urban, forest etc.) is still poorly known.

We surveyed spiders across an urbanization gradient in Guwahati city in the Indian state of Assam. We selected spiders as they are highly sensitive to environmental changes, particularly habitat modification and fragmentation (Miyashita et al. 1998; Bolger et al. 2000; Shochat et al. 2010). Attributes such as short life cycles, high abundances, conspicuousness and ecological sensitivity make spiders ideal focal organisms for this study (Kapoor 2008). Also, as different species exhibit different modes of hunting and habitat preferences, they can be classified into distinct guilds (Cardoso et al. 2011), which helps in assessing their functional diversity.

The specific objectives of our study were: (1) documenting the taxonomic richness of spiders in the study area, (2) understanding how their species diversity varies across habitats, and (3) understanding how their functional diversity changes across an urbanization gradient.

## Methodology

### Study site

The study was carried out in Guwahati (26°10’ N, 92°49’ E), the largest city of Assam state in northeast India. Guwahati city extends over 216 km² and supports a population of 1.12 million (Census of India 2011). The Guwahati Metropolitan Area (GMA), situated on the southern banks of River Brahmaputra, includes various hills, low-lying valleys, lakes and wetlands. It is flanked by the Buragosain Parbat in the east, and by the hills of Rani and Garbhanga in the south, contiguous with the hills of the neighbouring state of Meghalaya (Fig. 1). These hills constitute the major forest cover of the city. The city has undergone rapid urbanization in recent times, and it is among the top 20 developing ‘smart cities’ of India as per the Ministry of Urban Development, Government of India (Pawe and Saikia 2017). The geographic location of the city, between the banks of River Brahmaputra and the foothills of the Shillong Plateau, makes the city ideal for our study which involves documenting spider diversity across a gradient of habitats with different levels of urbanization.

**Fig 1:**
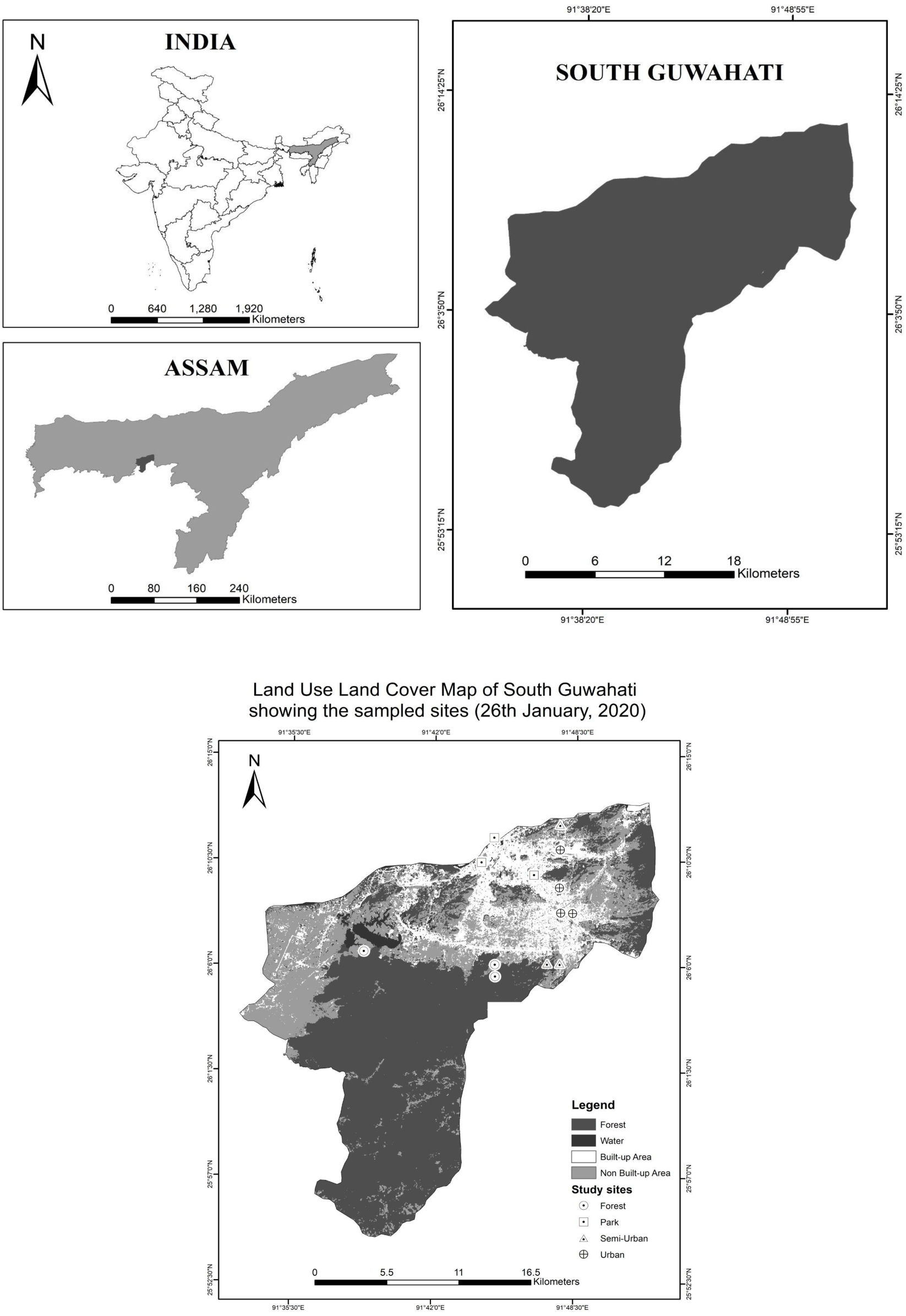
Land Use Land Cover Map of South Guwahati (2020)

Different spider species likely respond differently to changes in environmental conditions and geography (Parris et al. 2018). Hence, we identified five categories of habitat types based on the proportion of vegetation cover: urban (0–30%), public parks (30–40%), semi-urban (30–60%) and forest edge and core (> 80%). We used these habitat categories to select potential study sites based on the degree of urbanization determined using the land-use and land cover map of Guwahati (Fig 1). We used USGS Earth Explorer to acquire LANDSAT 8 satellite images and pre-processed them for atmospheric correction. We finally selected thirteen sites at random representing the five habitat types (Table 1).

**Table 1:** Study Sites and the species richness and diversity of spider species

| Name of the Sites | Site Identity | Habitat type | Species Richness | Species Diversity |
| --- | --- | --- | --- | --- |
| Shankar Dev Udyan, Bharalu | UP1 | Urban Parks | 24 | 0.76 |
| Shradhanjali Kanan | UP2 | Urban Parks | 21 | 0.84 |
| Nehru Park | UP3 | Urban Parks | 36 | 0.22 |
| Bamuni Moidam | U1 | Urban | 17 | 0.62 |
| Downtown | U2 | Urban | 17 | 0.36 |
| Hengrabari | U3 | Urban | 16 | 0.8 |
| Wireless | U4 | Urban | 7 | 0.7 |
| Basistha Forest Office | SU1 | Semi-Urban | 26 | 0.84 |
| IOC Noonmati | SU2 | Semi-Urban | 32 | 0.95 |
| Latakata | SU3 | Semi-Urban | 17 | 0.9 |
| Deepor Beel MD | FE1 | Forest Edge | 27 | 0.94 |
| Garbhanga Cave | FE2 | Forest Edge | 27 | 0.96 |
| Garbhanga Stream | FC1 | Forest Core | 27 | 0.95 |

### Sampling design

At each of the 13 selected sites, we established a 200 m long belt transect of width 10 m for surveying spiders. However, owing to constraints in access at certain sites, we had to limit the transect length to 150–200 m. All transects were surveyed for three days, except U1, UP1 and UP3, which were sampled for four days each. The various sampling days in each site constituted the temporal replicates, and the number of days of sampling was decided based on when we obtained a stable species accumulation curve for a particular site. Our study thus had a nested study design wherein transect surveys were ‘nested’ within sites that were nested within habitat types We conducted spider surveys from February 2020 through August 2020. Each transect survey was done over a 1–2 h period to ensure consistent sampling effort for each habitat type.

### Assessing spider diversity and abundance

We used a visual search sampling method for documenting species diversity and abundance. The advantage of this method is that spiders remain undisturbed and their numbers can be assessed repeatedly (Lubin 1978; Leite and Rocha 2022). During surveys, we carefully scanned all vegetation substrates such as tree bark, foliage, twigs, and branches and conducted ground searches under leaf litter and fallen/dry wood for spiders not visible directly on the ground. For identification, we used various published sources (e.g., Sebastian and Peter 2009; Assam State Biodiversity Board 2015) concerning spider diversity in India, particularly from Assam. We assessed the spider abundance as the number of spiders encountered per sample per site.

### Measuring functional diversity

Spiders have distinctive characteristics that facilitate the measurement of functional diversity. These include: foraging method, the range of prey hunted, vertical stratification, circadian activity, body size and phenology. We focused on three specific characters and 28 traits, including hunting mode (cursorial, ambush, foliage, cannibalism, foliage spitting, kleptoparasitism, web building and nocturnal hunting), type of webs they were found in (funnel, orb, nursery, sensing, sheet, silk retreats, space, trashline, tent, tangled, kleptoparasite found in the web of other species, jumpers found in silk or leaf sac, and no web) and preferred stratum (drain, grass, foliage, foliage over water, tree trunk, and wall/concrete surfaces). We selected these particular traits because they provide information about how the various habitats are utilized by spiders. Moreover, these trait types can be used to categorize different species into guilds.

### Data analysis

We considered the habitat type as the independent variable having five categories (urban, urban parks, semi-urban, forest core and forest edges), for species describing gradients in spider distribution. Mean abundance was calculated as the number of spiders per sample per site. The cumulative number of species found at each site represented species richness. We prepared species accumulation curves for each site to check the adequacy of our sampling efforts. We used the Simpson diversity index to quantify diversity as it accounts for both species richness and evenness. We used non-metric multidimensional scaling (NMDS) to identify groupings of similar sites based on a species composition matrix. We used the presence-absence data of various spiders in each of the 13 sites, to compute which sites were similar to each other in terms of species richness. We used the ‘ade4’ package in R to compute this analysis (Thioulouse et al. 2018). To further test whether there was a significant difference in spider species composition found across the various habitat sites, we computed the multi-response permutation procedure (MRPP) (McCune and Grace 2002). This analysis depends on the calculation of similarity indices calculated between pairs of sites (Shahabuddin and Kumar 2006). We used the Bray Curtis index for our data, which utilises species-wise differences by the total abundance in two plots (Ricotta and Podani 2017).

Further, we estimated the β-diversity to help determine whether the change in diversity, when we move from site to site or from habitat to habitat, is because of turnover or nestedness (Baselga 2010). We used ‘betapart’ package in R (Baselga and Orme 2012) R Core Team 2020) for this analysis, as it partitions the change in diversity across various habitats the turnover component, measured as Simpson dissimilarity (i.e., beta.SIM) and the nestedness component, measured as a nestedness-resultant fraction of Sorensen dissimilarity (i.e., beta.SNE).

We built generalized linear mixed models (GLMM) to identify which factor contributes the most to species richness and abundance. GLMM is an appropriate tool for analyzing complex ecological data because it includes random effects. Random effects can account for non-independence in data and inferences about the effects of fixed variables can be made more accurate and correct by modeling random effects (Harrison et al. 2018). We considered each of the 13 sites and their temporal replicates as random effects, while the habitat type provided for the fixed effect. Species richness and abundance were the response variables. We compared the richness estimates of semi-urban, urban, parks and forest edges taking the forest core as the baseline.

We used the FD package of R (Ver 1.0-12) for computing five functional diversity indices. These included functional dispersion (Laliberte and Legendre 2010), Rao’s quadratic entropy (Botta-Dukat 2005) and functional richness, functional evenness and functional divergence (Villeger et al. 2008). Functional richness (F_ric_) signifies the amount of functional space occupied by a given community (Villeger et al. 2008), while functional evenness (F_eve_) calculates the regularity of the abundance distribution in the functional space, and functional divergence (F_div_) measures how abundances tend to be on the outer margins of the functional space (Mouchet et al. 2010). Functional dispersion (F_dis_) is the mean distance of individual species to the centroid of all species in multidimensional trait space (Laliberte and Legendre 2010) and Rao’s quadratic entropy (RaoQ) measures the mean functional distance between two randomly chosen individuals (Mouchet et al. 2010). We conducted all the data analyses in R 4.0.0 and 4.0.2 (R Core Team, 2020).

## Results

We found 89 species of spiders belonging to 65 genera and 12 families. The suburban areas and urban parks recorded the highest number of species, followed by the forests and the urban areas. Forest core had the maximum diversity (D = 0.96) while urban parks had the lowest (D = 0.38), even though parks had more species (44) than forests (39). In terms of individual sites, the FE2 area had the highest diversity (D = 0.96) and U2 reported the lowest diversity (D = 0.36) (Table 1).

NMDS analysis revealed that three of the four urban sites grouped together, but U4 was dissimilar due to very low species richness. The park sites closely clustered with the three urban sites except for UP3 because the latter showed a much higher deviation in terms of species richness. Further, the distribution of semi-urban sites and forest sites was uneven due to greater dissimilarity in species composition (Fig 2). MRPP analysis of the spider species abundance–site matrix, suggested that the spider species composition differs significantly between the various habitat categories (P = 0.003).

**Fig 2:**
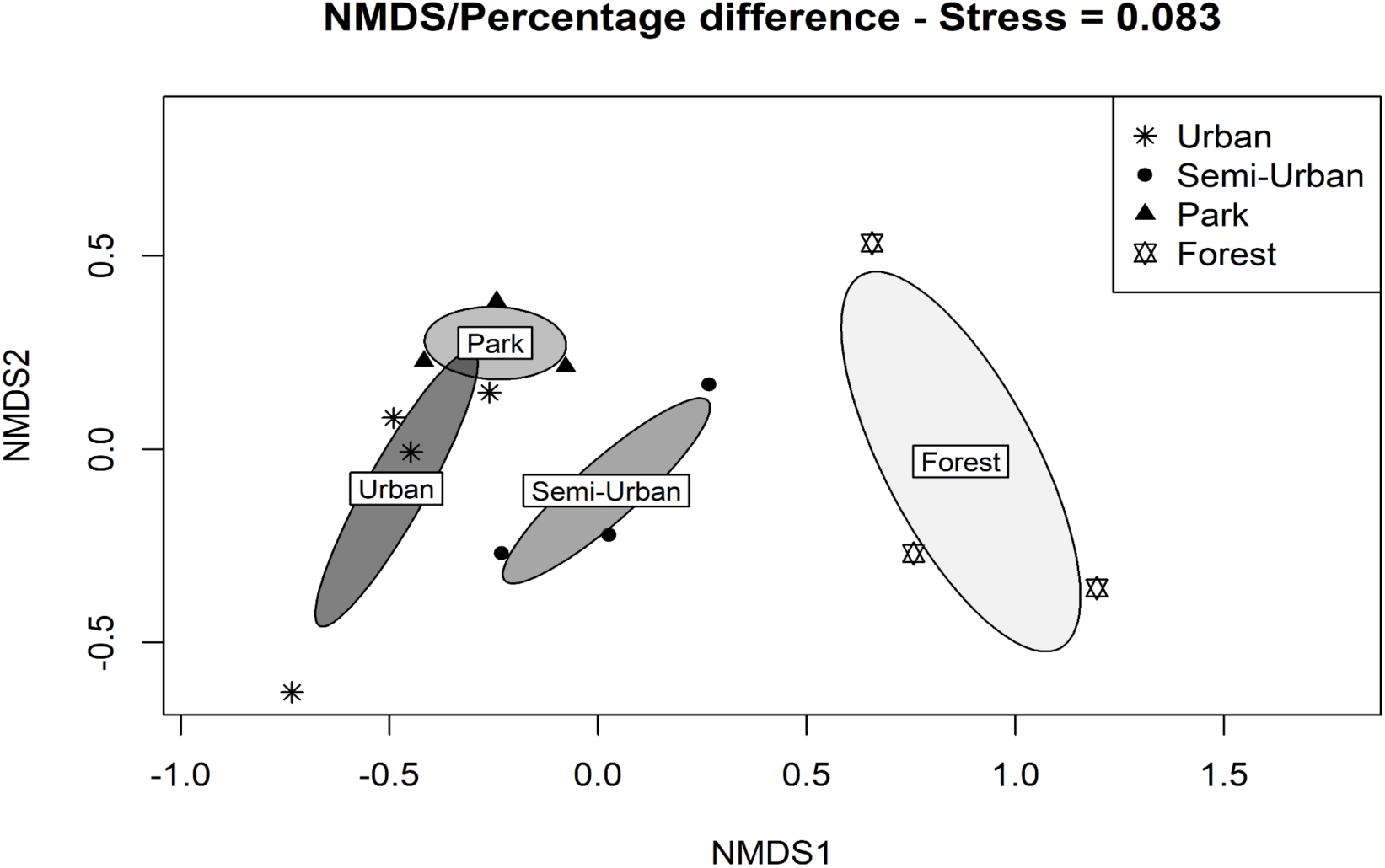
NMDS ordination of sites based on similarity in species composition

GLMM analysis showed no significant contributions of the random effects (sites and temporal replicates) on the species richness across the 13 sites (variance = 0.003 and 0 for sites and temporal replicates, respectively). Similarly, for abundance, the sites and temporal replicates had variances of 0.27 and 0.53, respectively.

The model with the habitat type (park, semi-urban, urban, forest edge and forest core) as a fixed effect exhibited contrasting results for both spider species richness and spider abundance across the 13 sites (see. Table 2 for summary). While the forest edge, park and semi-urban habitat types did not show deviation from the forest core, the richness in urban habitat was significantly different from that of the forest core.

**Table 2:** Summary of GLMM used to analyse the impact of habitat types (Forest core, Forest edge, park, semi-urban, urban) on the abundance and species richness of spiders across various sites

| Habitat Type | GLMM<br>Abundance<br>Estimate | Standard Error for<br>Abundance | GLMM<br>Richness<br>estimate | Standard Error<br>for Richness |
| --- | --- | --- | --- | --- |
| Forest core<br>(Intercept) | 0.61 | 0.390 | 2.71 | 0.112 |
| Forest edge | 0.69 | 0.674 | 2.8 | 0.189 |
| Park | 1.85 | 0.50 | 2.8 | 0.138 |
| Suburban | 1.12 | 0.502 | 2.75 | 0.143 |
| Urban | 1.77 | 0.476 | 2.18 | 0.148 |

The estimated abundance for the forest core was 0.61 and that of forest edge habitat was 0.08. Thus, the forest core and forest edge showed a negligible difference in abundance. However, parks, urban habitats and semi-urban habitat types significantly deviated from the forest core. The most pronounced deviations were in parks and urban sites.

The results from the ‘betapart’ analysis indicate that the contribution of turnover was greater (beta.SIM = 0.51) (Fig 3 (a)) than the contribution of nestedness (beta.SNE = 0.09) (Fig 3 (b)) in changing β-diversity as we moved from one habitat type to the other.

**Fig 3.**
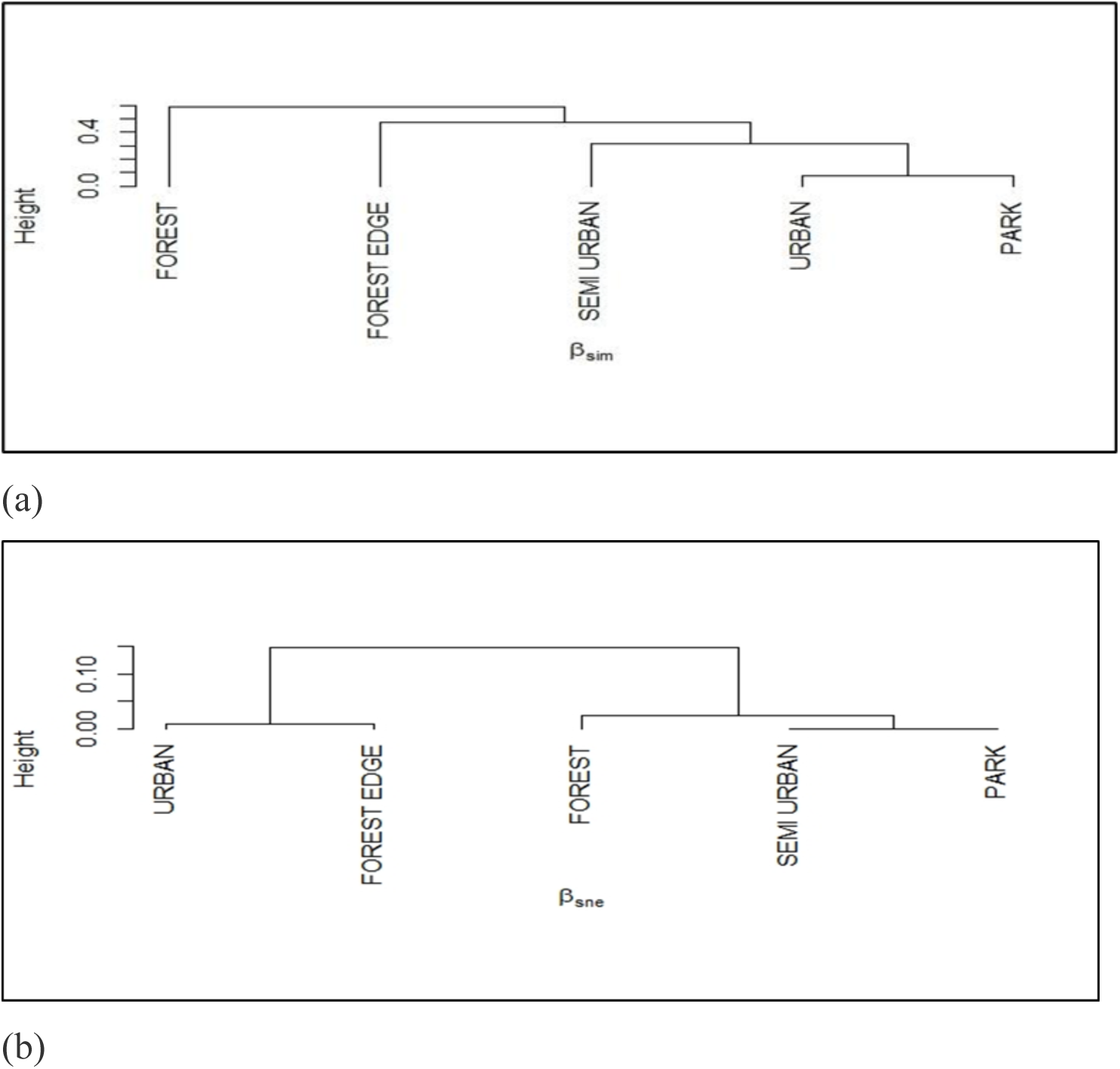
Dendogram of habitat types based on beta diversity contributed by (a) contributed by turnover (beta.SIM) and (b) nestedness (beta.SNE)

The functional diversity quantified the change in functional diversity parameters across three selected characters and 28 different traits (Table 3). The functional richness was highest in forest core (F_ric_ = 23.43) and lowest in urban habitats (F_ric_ = 12.98). The functional divergence, on the other hand, showed an opposite trend where the urban areas had the greater trait divergence (F_div_ = 0.79) as compared to the forest core (F_div_ = 0.78). Forest edge showed the highest evenness in the distribution of traits as well as in trait dispersal (F_eve_ = 0.48 and F_dis_ = 1.83). In terms of individual sites, FE2 represented the maximum functional richness (F_ric_ = 0.96) and the minimum was represented by UP3 (F_ric_ = 0.22).

**Table 3.** Habitat-wise data on functional diversity parameters

| Habitat type | Number of Species | Diversity | Functional richness ($F_{ric}$) | Functional evenness ($F_{eve}$) | Functional diversity ($F_{div}$) | Functional dispersal ($F_{div}$) | Rao's quadratic entropy (RaoQ) |
| --- | --- | --- | --- | --- | --- | --- | --- |
| Sub-urban | 44 | 0.91 | 16.68 | 0.34 | 0.78 | 1.27 | 2.43 |
| Forest core | 39 | 0.96 | 23.43 | 0.45 | 0.74 | 1.47 | 2.65 |
| Forest edge | 28 | 0.94 | 21.40 | 0.48 | 0.71 | 1.83 | 4.05 |
| Urban | 26 | 0.54 | 12.98 | 0.36 | 0.79 | 1.12 | 1.94 |
| Park | 44 | 0.38 | 22.70 | 0.36 | 0.72 | 0.55 | 0.91 |

## Discussion

Our study shows that both the taxonomic and functional diversity were highest in forests and lowest in parks and urban sites, respectively. A similar pattern was observed in for functional richness and evenness. Conversely, the maximum trait divergence was found in urban areas. Turnover or replacement of species, possibly due to habitat differences, was the major contributor to β-diversity changes. Moreover, the results from the GLMM analysis suggested that the habitat types had a significant impact on the abundance in each site. Observations concerning the traits indicated that most ambush hunters were found only across highly vegetated sites like parks, semi-urban and forest sites and only a few were found in urban concrete regions. Web hunters were the dominant groups in the urban residential sites. No specific pattern was seen in the distribution of ground-dwelling jumpers.

As suggested by the observed results, the taxonomic diversity of spiders declined as we moved from an urban environment to semi-urban sites and finally to forest habitats (Table 1). There was much greater evenness in the distribution of species in the forest core and edge. Contrastingly, even though parks had the highest species richness, the habitat type marked the dominance of a single synanthropic species of spider *Cyrtophora cicatrosa* whose abundance was 100–300, whereas the other species had an abundance of only 1–20 individuals per species. Synantrophy is the temporal and spatial response of various species in response to urbanization (Guetté et al. 2017). Even in the case of semi-urban areas, the dominance of a few common spiders was responsible for lower species diversity as compared to forests.

Biotic homogenisation is the process by which two or more spatially distributed ecological communities become increasingly similar over time (Olden et al. 2018). The dominance of synanthropic species in urban habitats can be because of local disturbances caused by anthropogenic activities that create such a homogenization gradient, in which synanthropic species are found at the urban core, whereas urban fringe habitats and suburban areas comprise successional plants and edge animal species which are mostly native (McKinney 2006; Lowe et al. 2018). This can be because of the availability of mixed vegetation and a greater amount of leaf litter in these habitat types (Batery et al. 2018; Lowe et al. 2018) Additionally, the lack of vegetation cover, large concrete surface and regular maintenance and cleaning of private lawns or drainage as observed during our sampling period, could have been some of the attributing factors responsible for the very low species richness and diversity of spiders in the urban area as compared to the other habitat types.

The random variables in the present study i.e., each of the 13 sites and the temporal replicates did not have any significant impact on species richness or abundance. However, the modelling of fixed effect showed that there was a marked difference in the abundance of species in parks, semi-urban areas and urban areas with respect to forest core. This was again because the abundance in these three habitat types was much higher due to the presence of a few dominant species. This is in contrast to other studies where it was found that urbanization has a negative impact on abundance (Marzluff 2001; Buczkowski and Richmond 2012; Scheffers and Paszkowski 2012; Piano et al. 2020). Moreover, since the richness in forest core, forest edge, semi-urban and park habitats were relatively similar therefore we can conclude that urban habitats were also significantly different from the forest edge, park and semi-urban habitats in terms of spider richness.

In order to understand the underlying processes of biodiversity and their causes, we need to assign respective biological patterns to different β-diversity (Baselga 2010). Beta diversity analysis partitions the biological patterns into two components-turnover, species replacement and nestedness, or species loss. The change in the diversity of spiders between different habitat types is majorly due to turnover, indicating species replacement because of dispersal or niche processes. This implies that all the habitat types contributed more or less equally to regional diversity over time. Estimating the contribution of the various components of beta diversity is important in terms of conservation because a dominating turnover suggests the need to prioritize a regional approach of conservation focusing on multiple sites, whereas a dominating nestedness will shift the conservation priority to a few high diversity sites (Angeler 2013).

Changes in functional diversity can influence various ecosystem processes like ecosystem dynamics and nutrient availability (Goswami et al. 2017). Changes in the environment due to anthropogenic activities can also affect the functional diversity of the local population. In fact, studies have stated that increased urban activities are responsible for the reduction in functional diversity (Carrero et al. 2009; La Sorte et al. 2018; Tóth and Hornung 2020). Urbanization can be, hence, termed as human-induced rapid environmental change (Dahirel et al. 2017) that causes a variation in the adaptive behaviours of the organisms in the ecosystem. The observed tendency of a decreasing functional richness with increasing urbanization may be because of niche filtering caused by high intensities of land use in less-heterogeneous vegetation types that result in the clustering of communities with reduced ecological specialization (Wong et al. 2019). In contrast, the availability of a number of heterogeneous and complex niches in forests reduces the effect of land use intensities on the functional structure of organisms (Wong et al. 2019). However, the increased divergence in traits in urban areas, as obtained in our study, contradicts this assumption of niche filtering and hence puts forward scope for further research and understanding of this parameter of functional diversity measurement. Understanding the impact of land use changes due to urbanization is important for maintaining functional diversity as well as for conservation practices (Vandewalle et al. 2010). In case of species loss due to urban-induced disturbances, for example, functional diversity will be maintained even in disturbed environments, if the lost species represent similar traits to the remaining ones (Matuoka et al. 2020).

## Conclusion

The role played by urbanization in changing the biodiversity of a region is crucial in understanding the anthropogenic causes of changing dynamics of a community or an ecosystem as a whole. Moreover, it also provides scope for quantifying the impact of habitat fragmentation. Our study reveals that urbanization has a negative impact on the taxonomic as well as functional diversity of spiders. This result calls attention to effective conservation practices within urban cities as urban vegetation patches have the potential to shelter high species richness and abundance. In fact, the incorporation of vegetation patches within the urban environment, followed by proper maintenance of these patches and reducing disturbance at the microhabitat level can perhaps help in maintaining the arthropod diversity. Moreover, the dominating trend of turnover from the study makes it easier for us to prioritize the conservation of spiders at a regional level, as each site harbour a unique range of species.

The impact of urbanization on functional diversity, however, is still in the need of extensive exploration, in order to come up with a concrete trend or theory. Nevertheless, it is noteworthy that functional diversity being a relatively novel subject provides opportunities for further research in understanding the role of anthropogenic activities in modifying the functioning of an ecosystem.

## Acknowledgements

We would like to extend our deepest sense of gratitude towards Swagata Devi, Dhiraj Kumar Das and Ananya Bora for their assistance and guidance throughout the project. We would also like to thank the forest officials of Garbhanga Reserved Forest and the staff of Medicinal and Aromatic Plants Garden, City Plantation range Chakardo, for their help and cooperation.

